# Increasing temporal variance leads to stable species range limits

**DOI:** 10.1101/2021.08.09.455156

**Authors:** John W. Benning, Ruth A. Hufbauer, Christopher Weiss-Lehman

## Abstract

What prevents populations of a species from adapting to the novel environments outside the species’ geographic distribution? Previous models highlighted how gene flow across spatial environmental gradients determines species expansion vs. extinction and the location of species range limits. However, space is only one of two axes of environmental variation — environments also vary in time, and we know temporal environmental variation has important consequences for population demography and evolution. We used analytical and individual-based evolutionary models to explore how temporal variation in environmental conditions influences the spread of populations across a spatial environmental gradient. We find that temporal variation greatly alters our predictions for range dynamics compared to temporally static environments. When temporal variance is equal across the landscape, the fate of species (expansion vs. extinction) is determined by the interaction between the degree of temporal autocorrelation in environmental fluctuations and the steepness of the spatial environmental gradient. When the magnitude of temporal variance changes across the landscape, stable range limits form where this variance increases maladaptation sufficiently to prevent local persistence. These results illustrate the pivotal influence of temporal variation on the likelihood of populations colonizing novel habitats and the location of species range limits.

## Introduction

The limits of species’ geographic distributions have long puzzled biologists [1,2]. The causes of distribution, or range, limits fit into two broad categories — either populations have not colonized suitable areas outside their current range margin (a distribution limited by dispersal) or the environment outside that margin is unsuitable, keeping population growth below replacement (a distribution limited by adaptation). The latter case begs the question of why populations do not simply adapt to the novel environmental conditions beyond their range edge. This question is especially perplexing as this exact process — adaptation to novel environments — adaptation to novel environments adaptation to novel environments — presumably gave rise to the species’ *current* distribution.

Spatial gradients in the environment clearly play a major role in determining range limits — as the environment changes across space, so do the observed flora and fauna. Climbing a mountain slope, one can readily observe how community composition changes with elevation. But *within* species, there is ample evidence for populations successfully adapting to all sorts of abiotic and biotic environmental gradients [3–8]. What causes this adaptive process to cease and a range limit to form? Theory has shed light on the mechanisms underlying the relationship between spatial environmental gradients, local adaptation, and species distributions. For a species occupying a landscape with a spatial gradient (e.g., in soil pH, precipitation, etc.), theory tells us that several mechanisms may constrain adaptation, and subsequently, expansion at the edge of a range. Steep environmental gradients and high gene flow can swamp adaptation at the range edge to create stable range limits [9,but see 10]. Metapopulation dynamics [11] and biotic interactions [12,13] also can enforce range limits. Recent simulation approaches have highlighted how demography, genetic drift, expansion load, and not only the slope, but the shape (e.g., linear vs. non-linear) of the environmental gradient influence adaptation, range expansion, and the formation of stable range limits [14–17]. In all of these theoretical treatments, the spatial environmental gradient, which is modeled as a spatial gradient in phenotypic optima, is key to understanding when populations can expand and when stable range limits form.

However, habitats vary not only in space, but also in time. Indeed, in nature, temporal variation in the environment is the rule rather than the exception [18–20]. Variation in weather within and between years provides the most obvious illustration of abiotic temporal variation. Biotic environments also fluctuate (stochastically or regularly) through time as populations of predators, mutualists, pathogens, and competitors wax and wane. Both theoretical and empirical work demonstrate that temporal variation has important consequences for population demography [21–24,reviewed in 25] and evolution [20,26–31]. Demographic models have illustrated that a population’s extinction risk generally increases as temporal variation increases [23,32], and experimental work has supported these theoretical predictions [24,33]. Viewed through an evolutionary lens, temporal variation can generate fluctuating selection due to phenotypic optima changing through time [34]. These changes in phenotypic optima can lower expected individual fitness via increases in genetic load, i.e., the reduction in fitness due to an individual’s deviation from an optimal phenotype [35]. With phenotypic optima changing through time, even a phenotype perfectly adapted to the long-term mean optimum will experience fitness costs due to genetic load [36,37]. In microcosm experiments, temporally fluctuating environments have also been shown to impede adaptation to directional environmental changes, presumably due to relaxed selection during benign periods [30]. However, Holt et al. [38] showed how temporal environmental fluctuations can have positive effects and facilitate adaptation in sink habitats if the sink environment becomes more benign long enough to increase population growth and adaptation. Similarly positive effects of temporal variation were observed in an experiment exposing diatoms to increasing temperatures [31], where increased population sizes during benign periods resulted in more effective selection. Though our understanding of the prevalence of fluctuating selection in natural populations is still incomplete and plagued by sampling error [39,40], several recent rigorous studies do show strong temporal fluctuations in selection [e.g., 34,41,42], implying fluctuating phenotypic optima through time.

Because temporal variation can influence key aspects of population demography and evolution, there is reason to expect it influences species range dynamics, as well [43]. However, most range limit models to date assume temporally constant environments. Yet in nature, the environment will always vary in both space and time, and it is easy to imagine myriad ways that temporal variation might affect the fate of populations spreading across a spatial gradient. If the environment is changing directionally through time (e.g., glacial advance, warming temperatures), theory has shown how the rate of environmental change, the amount of genetic variance, and the steepness of an underlying spatial gradient all influence population persistence [44,45]. Recent work by Holt et al. [43] provided some of the first insights into the influence of non-directional temporal variation on range dynamics, showing how temporal variation in competition can modulate the size of an established species’ range, and that temporal variation in immigration increases the probability of establishment in sink habitats. However, we lack range limit models exploring the spread of a species across continuous spatial gradients with non-directional temporal variation in environmental conditions. Such temporal variation in the environment could slow adaptation along a spatial gradient via fluctuating selection. Alternatively, positive demographic effects of temporal variation, such as high fecundity in a relatively favorable year, could boost population sizes and increase range expansion and the efficacy of natural selection [38]. Temporal variation will directly affect extinction probabilities via demography [23], and could further influence adaptation by influencing levels of genetic variation [46]. Importantly, just as environmental variation through space can manifest with different patterns (e.g., linear / non-linear gradients), environments can vary through time in different ways. First, environmental variation can exhibit temporal autocorrelation patterns that are negative (e.g. dry years tend to be followed by wet years and vice versa), positive (e.g. dry years tend to be followed by another dry year), or reflect uncorrelated random noise [47]. Second, temporal variance may not be equal across space. For instance, positive relationships between demographic variability and distance from the center of a species’ range suggests that habitats on the edge of the range may sometimes be more temporally variable than habitats in the core [48,49].

In this paper, we use analytical and simulation models to ask how temporal variation in the environment influences population expansion and the formation of range limits across spatial gradients. In our models, temporal variation manifests as temporal stochasticity in phenotypic optima, with optima fluctuating generation to generation around the patch’s long-term mean optimum. We explore two geographic modes of temporal variation (Fig. 1):

**Figure 1.**
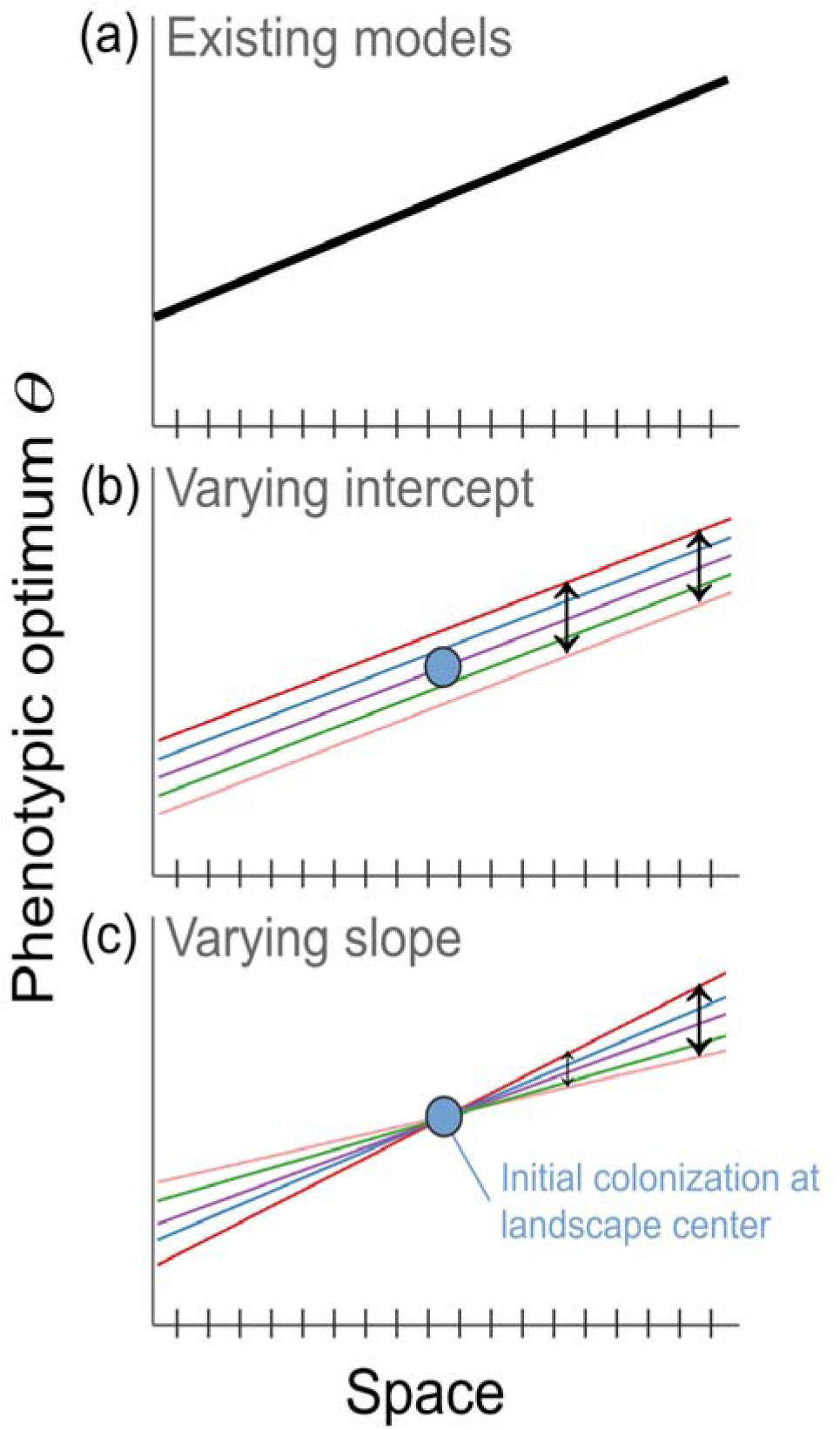
Conceptual diagram illustrating how temporal variation is incorporated in the model. (a) illustrates how spatial environmental variation is modeled in existing range limit models, with a temporally constant spatial gradient in phenotypic optima. In our model, there is intergenerational change in phenotypic optima around the mean optimum at each point in space, according to two “geographic modes” of temporal variation. In the “varying intercept” scenario (b), the intercept of the spatial phenotypic optimum gradient fluctuates such that all patches experience the same magnitude and direction of deviation from the mean optimum each generation. In the “varying slope” scenario (c), the slope of the spatial gradient fluctuates stochastically around the mean slope each generation. The gradient is “anchored” at the center of the landscape, thus, the magnitude of temporal variance increases with distance from the landscape center. The blue dots (b, c) represent the initial founding population in the simulation, which then spreads (or does not spread) across the landscape. Black arrows indicate the magnitude of temporal variance at that point on the landscape [equal at all points in “varying intercept” and increasing with distance from the landscape center in “varying slope”].

1. **Varying intercept**: all points in space experience environmental change equally through time. This would be the case if, say, temperatures or precipitation changed across a landscape by a similar magnitude and direction year to year (e.g., a landscape-wide drought). It manifests as stochasticity in the intercept of the spatial gradient in phenotypic optima, fluctuating around the long-term mean spatial gradient (Fig. 1b)
2. **Varying slope**: the magnitude of environmental change through time increases away from the landscape center. This would be the case if, say, there were increased temporal variation in inundation at the edge of a wetland compared to the center, or increased temporal variation in rainfall as one goes from mesic to arid habitat. It manifests as stochasticity in the slope of the spatial gradient in phenotypic optima, with a constant intercept at the range center. Thus, the phenotypic optimum displays larger fluctuations around the long-term average with increasing distance from the range center (Fig. 1c).

We first derive a simple analytical model to describe how, under these two different geographic modes, temporal variation imposes genetic load on populations across a landscape. We then explore the influence of temporal environmental variation on range expansion and dynamics using an individual-based, forward-in-time genetic simulation model. In the model, a single population colonizes the center of a spatial environmental gradient, and we track the demography, evolution, and spread of populations across the landscape with the environment varying in space and time. We vary both the steepness of the underlying spatial environmental gradient and the geographic mode of temporal variation as defined above. We also vary the temporal correlation pattern of the environment from negative to positive autocorrelation.

## Materials and Methods

### Data Availability Statement

All SLiM and R code, simulation parameters, and the simulation results needed to reproduce the figures in this manuscript, are uploaded for review and will be made available on Figshare upon acceptance.

### Expected genetic load due to temporal variation

We first present a simple analytical model to describe how our two geographic modes of temporal environmental variation (varying intercept and varying slope) affect expected individual fitness. Consider a species distributed across a landscape, with populations occupying discrete patches. The environment varies across space (*x*), producing a spatial gradient in phenotypic optima with slope *b* (Fig. 1). Each population comprises individuals perfectly adapted to the long-term mean phenotypic optimum at that point (patch *x*) in space. As the environment varies stochastically through time, mismatches arise between individuals’ phenotypes and the current phenotypic optimum, even though these individuals are perfectly adapted to the long-term mean phenotypic optimum within their patch. We model this mismatch as genetic load, which is the fitness cost incurred by an individual due to deviation from the phenotypic optimum [35]. As in Lande and Shannon [36], we define genetic load for an individual *i* as

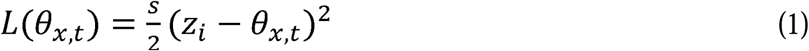

where *θ*_*x,t*_ is the phenotypic optimum in patch *x* at time *t, s* is the strength of stabilizing selection, and *z*_*i*_ is the individual’s phenotype. In our varying intercept scenario (Fig. 1b), *θ*_*x,t*_ = *ε*_*t*_ + *bx*, while in our varying slope scenario (Fig. 1c), *θ*_*x,t*_ = (*ε*_*t*_ + *b*) *x* In both cases, *b* is the slope of the gradient in phenotypic optima, *x* is the patch position along the gradient, and *ε*_*t*_ is a normally distributed random variable defining the stochastic deviation in either the intercept or slope of the spatial environmental gradient.. For now, we assume uncorrelated temporal fluctuations with a fixed variance of *τ* (i.e., *ε*_*t*_ ∼ *N*(0, *τ*)). For an individual perfectly adapted to the expected phenotypic optimum (i.e., *z*_*i*_ = *bx*), genetic load in the varying intercept scenario becomes

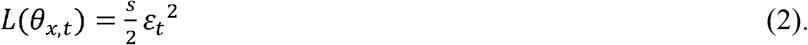

Following mathematical rules for transformations of normally distributed random variables yields

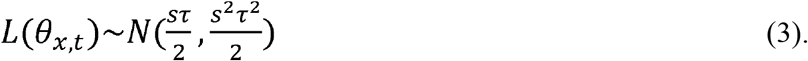

Similarly, genetic load for an individual with *z*_*i*_ = *bx* in the varying slope scenario is

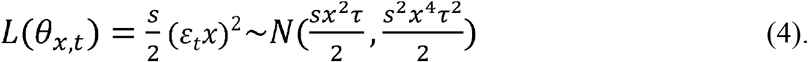

Therefore, the mean and variance in genetic load increase with spatial position (*x*) in the varying slope scenario, but are constant in space in the varying intercept scenario.

### Effects of temporal variation on range dynamics

The analytical model above describes individuals’ expected genetic load due to temporal stochasticity in phenotypic optima in our two landscape scenarios. However, the model does not account for gene flow, density dependence, genetic drift, demographic stochasticity, varying patterns of temporal autocorrelation, or range colonization dynamics. To explore these factors, we used SLiM [50], a forward-in-time population genetic modeling software, to build a complex, individual-based simulation of species range dynamics in a spatially and temporally varying environment. SLiM has recently come to the fore as a flexible, fast, and powerful tool to model individual genomes under a wide variety of spatially and temporally explicit evolutionary scenarios. Our simulation model builds most directly on the modeling frameworks of Polechová and Barton [14] and Bridle et al. [16], which explored the role of spatial environmental gradients on range dynamics. We build on the insights from those models by asking how temporal variation in the environment influences population spread and the formation of range limits.

### Genetics and mutation

In our simulations, individuals were diploid and either male or female, with obligate sexual reproduction, and a single chromosome 100,000 bp long. There were two mutation types: 1) neutral mutations, and 2) mutations that contributed additively to a quantitative trait [i.e., biallelic quantitative trait loci (QTL) with no dominance]. The overall mutation rate was set to 1 ⍰ 10^−7^ mutations per base position on a gamete per generation (Table S1), and mutations were 10 times more likely to be neutral than QTLs. QTL effect size was drawn from a standard normal distribution (i.e. N(0, 1)). Recombination rate was set to 1 ⍰ 10^−5^ (probability of a cross-over event between any two adjacent bases per genome per generation).

### Mating and population dynamics

The simulated landscape comprised a one dimensional array of 201 patches. This is most analogous to a natural landscape that approximates one dimension, like a river, river corridor, mountain ridge, or valley. Each patch hosted a local population subject to density-dependent regulation, with carrying capacity (for a perfectly adapted population) constant across the landscape (here, K = 50). Following Bridle et al. [16], individual fitness (*W*_*i*_) was calculated as

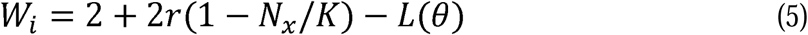

Where *r* is the population maximum rate of increase (set to *r* = 0.8; Table S1), *N*_*x*_ is the number of individuals in population *x, K* is the carrying capacity of each patch (set to *K* = 50), and *L*(*θ*) defines the genetic load as defined in the analytical model above (Eq. 1). Thus, the first part of equation (5) describes standard logistic growth and the second part introduces the fitness cost scaled by the deviation of an individual’s phenotypic value from the local optimum (i.e., genetic load). For females, *W*_*i*_ defined the mean of a Poisson distribution for the number of offspring that female produced (female fecundity); for males, *W*_*i*_ defined the likelihood of a male mating (*W*_*i*_ corresponded to a male’s weight in a weighted draw from the pool of available males in a population). Males could mate multiple times. Any *W*_*i*_ < 0 was set to *W*_*i*_ = 0.00001 (as the mean for a Poisson distribution in SLiM must be greater than zero). After mating occurred within a population, offspring dispersed according to a Poisson dispersal kernel with mean (*m*) = 0.8. The direction of dispersal (left or right along the gradient) was unbiased and random. Individuals set to disperse beyond the edges of the landscape were instead routed to the most distal patch in that direction. To avoid any potential edge effects, one end condition for our simulations (see below) was if the population of either of the most distal patches reached half the carrying capacity or more. It is important to note that dispersal (*m*) and carrying capacity (*K*) are both hard to estimate and highly variable in natural populations; for our main simulations we use values for these parameters near the mode of their distributions as estimated by Polechová and Barton [14]. We explored the sensitivity of our results to varying *m* and *K* in Supplementary Material S1.

### Environmental variation

Environmental variation manifested as changes in the optimum phenotype of the quantitative trait across space and time. The slope of the spatial environmental gradient (*b*) represents the change in phenotypic optima across the landscape (Fig. 1a). To illustrate how *b* manifests as a fitness cost of dispersal (i.e., genetic load due to spatial variation in optima), let us assume *s* = 0.125 (used throughout; Table S1). Then the genetic load incurred by dispersal to patch *x+1* for an individual perfectly adapted to patch *x* is,

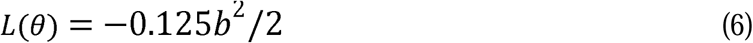

Thus if *b* = 1, a female perfectly adapted to patch *x* will experience a fitness cost of *L*(*θ*) = 0.0625 upon migration to patch *x+1*, or a fitness decrease of ∼3.125%.

Temporal environmental variation was implemented as intergenerational stochasticity in the intercept or slope of the spatial gradient in patch phenotypic optima using the same equations presented for the analytical model above. To incorporate temporal autocorrelation, the random deviations in intercept or slope were defined as:

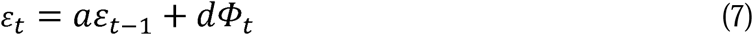

where *ε*_0_ = 0, *d* scales the magnitude of noise *Φ*_*t*_ ∼ *Normal*(*0, τ*), and *a* represents the degree of temporal autocorrelation. When *a* = 0 and *d* = 1, Equation 7 is equivalent to the distribution of stochastic deviations in phenotypic optima used in the analytical model above. Temporal autocorrelation could be positive (0 < *a* ≤ 1), negative (−0.99 ≤ *a* < 0), or uncorrelated (*a* = 0) temporal stochasticity, with autocorrelation lag equal to one generation (Fig. S1). (We prevented *a* from going all the way to −1 because, due to *ε*_0_ = 0, this would be identical to *a* = 1.) We set *d* = (1 − *a*^2^)^0.5^ so that *Var*(*ε*) over the full time series used in our simulations was approximately equal for all values of *a*.

*τ* differed between the two geographic modes of temporal variation. For the “varying intercept” scenario we set *τ*_*intercept*_ = 4. This resulted in temporal variation in selection gradients, quantified as *σ*_*β*_ ≈ 0.14, being within the range of estimates from natural populations (median *σ*_*β*_ = 0.099 in de Villemeruil et al. [42], averaging across birds and mammals; full derivation of *σ*_*β*_ presented in Supp mat. S2). For the “varying slope” scenario, we set *τ*_*slope*_ = 0.0025 which resulted in patch conditions mimicking the “varying intercept” scenario at 40 patches away from the landscape center (Fig. 2). We also ran simulations with the intercept and slope varying simultaneously; in this “varying intercept and slope” scenario, respective *τ* values remained the same (*τ*_*intercept*_ = 4 and *τ*_*slope*_ = 0.0025).

**Figure 2.**
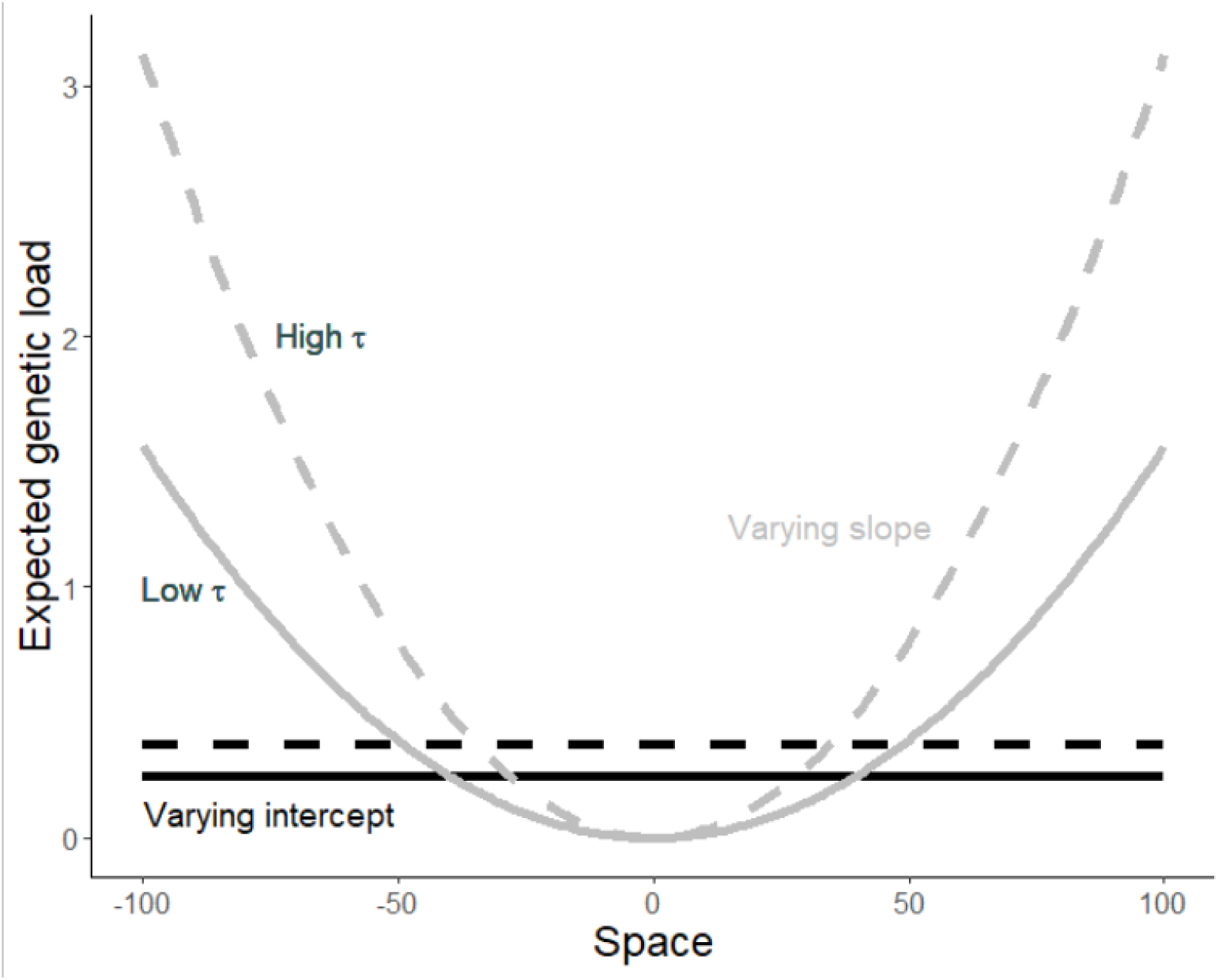
Results of the analytical model estimating expected genetic load across space due to temporal stochasticity in phenotypic optima, for the varying intercept (black; Fig. 1b) and varying slope (gray; Fig. 1c) scenarios. Solid lines represent lower magnitude of temporal variance (and); dashed lines represent higher magnitude of temporal variance (and)

### Simulation process

#### Burn-in

Each simulation began with a burn-in period of 20,000 generations. At the start of the burn-in, 100 genetically homogenous, perfectly adapted, founding individuals were initiated in the central patch of the landscape. Mating and dispersal happened as described above, but the landscape was limited to 21 patches wide (10 patches on either side of the founding patch). Carrying capacity in each patch was 100 individuals (thus, landscape-wide K = 2,100). There was a modest spatial environmental gradient in optima across these 21 patches (b = 0.5), and modest, uncorrelated temporal fluctuations in the intercept of the spatial gradient in optima [deviations drawn from Normal(0,1)]. The goal of this burn-in period was to generate independent replicates of genetically variable source populations that had evolved in landscapes of similar spatial and temporal heterogeneity for each simulation run, while allowing enough time for the different simulation burnins to converge on similar levels of genetic diversity. During the burn-in, mean heterozygosity of neutral mutations (*π*) in the central population usually reached an equilibrium by 10,000 generations (Fig. S2). At generation 20,000, a random subset of 50 individuals was selected from the 21-patch landscape and migrated to the central patch for the “founding event”. All other individuals were then removed from the simulation.

#### Main simulation

After the burn-in period, the main simulation began with the prescribed parameters and the available landscape expanded to 201 patches, with 50 individuals in the founding central patch. Simulations ended after 20,000 generations, or if all populations went extinct, or if at least one of the most peripheral landscape patches reached a population size at least half the carrying capacity (i.e., the species had filled the entire landscape). We ran 1000 simulations for each of the three temporal variation scenarios (varying intercept, varying slope, varying intercept and slope) with parameter values pulled randomly from uniform distributions: *b*[0-3], *a*_*intercept*_[-0.99-1], *a*_*slope*_[-0.99-1]. To compare our model with the model of Polechová & Barton [14], we also ran 200 simulations, varying *b* but with *a*_*intercept*_ = 1 and *a*_*slope*_ = 1 (i.e., a temporally constant environment), and compared our results to results predicted from that model (Supp. mat. S3).

## Results

### Analytical model

As previous work has shown [29,36,38], temporal variation in phenotypic optima results in a genetic load on individuals (Fig. 2). In our varying intercept scenario, this load is constant across space, while in the varying slope scenario, load increases non-linearly with distance from the landscape center. Genetic load increases with the magnitude of temporal variance () and the strength of stabilizing selection (*s*).

### Simulation model

With a temporally constant environment (i.e., *a* = 1) expansion was prevented where the spatial gradient slope (*b*) was ⍰ 2.5, somewhat steeper than the threshold predicted from the model of Polechová & Barton [14], *b* 1.96 (Supp. mat. S3).

### Varying intercept

When temporal variation in the environment was equal in all patches on the landscape (“varying intercept” scenario; Fig. 1b), a clear relationship emerged between the degree of temporal autocorrelation (*a*) and the slope of the underlying spatial gradient (*b*) in determining whether populations spread across the landscape or went extinct (Fig. 3). Range expansion was favored when temporal variation was more positively autocorrelated and spatial gradients were shallow. The ultimate fate of the species in each simulation was either eventual extinction or continual expansion; stable range limits did not form, though the rate of range expansion slowed as spatial gradients steepened (light blue points in Fig. 3). (Here and below we use “species” to describe the group of populations on the landscape.)

**Figure 3.**
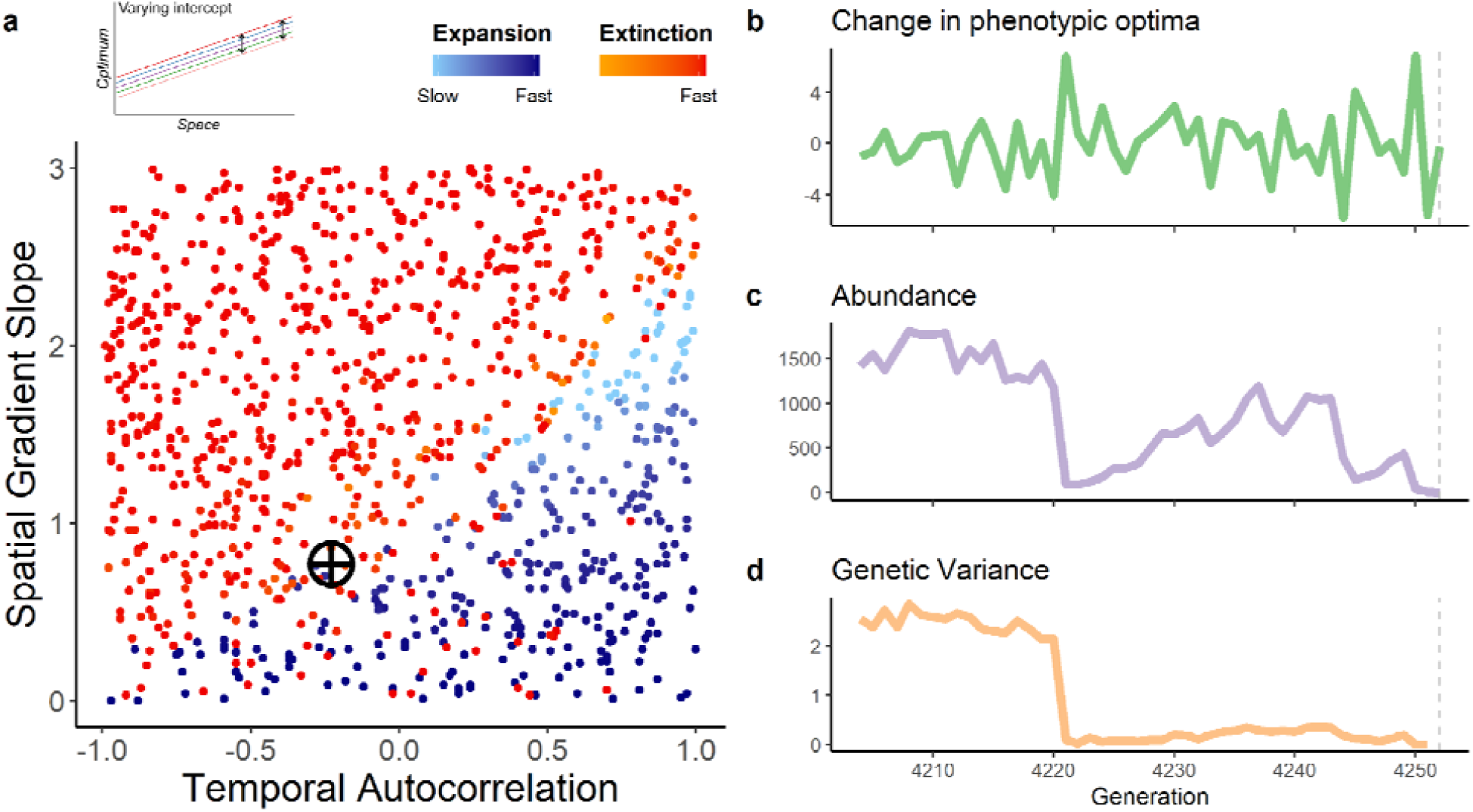
Range dynamics under the “varying intercept” scenario. (a) When temporal variation manifests identically across the landscape, range expansion is favored when temporal variation (*a*, X-axis) is more positively autocorrelated and spatial gradients (*b*, Y-axis) are shallow. (A temporal autocorrelation of 1 is a temporally static environment, as modeled in previous range limit models.) Each point represents a simulation (N = 1000). No stable range limits formed in this scenario; species either continually expanded or went extinct. All blue points are simulations where species were able to spread, and red/orange points are simulations where species went extinct; points are colored by how quickly they expanded or went extinct. (b-d) Details of one simulation (marked with crosshairs in panel a) prior to extinction. Panels track (b) temporal changes in patch phenotypic optima, (c) landscape-wide population abundance, and (d) median population genetic variance (for the trait conferring adaptation to the gradient) across generations prior to extinction (grey dashed line).

As in previous models, steep spatial gradients introduced a strong fitness cost of dispersal, allowing drift to overpower selection and stymie adaptation. Temporal variation exacerbated the negative effects of dispersal by introducing a fitness cost even for individuals well-adapted to their patch’s long-term mean optimum; i.e., temporal fluctuations in optima meant that no genotype could completely escape maladaptation across generations. In environments with temporal variation, populations were often able to adapt to the underlying spatial gradient if it was not too steep. However, a large deviation from the mean optimum in a generation had large negative demographic consequences due to individuals being strongly maladapted in most patches. If then the following generation experienced a strongly different optimum, which is more likely as temporal autocorrelation becomes more negative, extinction risk was high (Fig. 3b,c). Thus, temporal variation introduced an extinction risk due to fluctuating phenotypic optima that had strong effects on mean fitness [e.g., 29,36,38].

### Varying slope

When the magnitude of temporal variance increased away from the landscape center (“varying slope” scenario), stable range limits formed (Fig. 4). The exact location of the range limit fluctuated over time as extinction/colonization dynamics played out at the range edge; thus, we define stable range limits as the most distal patches on either side of the founding patch that did not go extinct for at least 950 of the 1000 generations before the end of the simulation (which lasted 20,000 generations). Range limits formed where increasing temporal variance resulted in increasing genetic load (Fig. 2), causing populations to fail to adapt to the optimum of edge patches because temporal fluctuations in optima caused populations in those patches to often go extinct (or close enough to extinction to strongly undermine genetic variance and adaptive potential).

**Figure 4.**
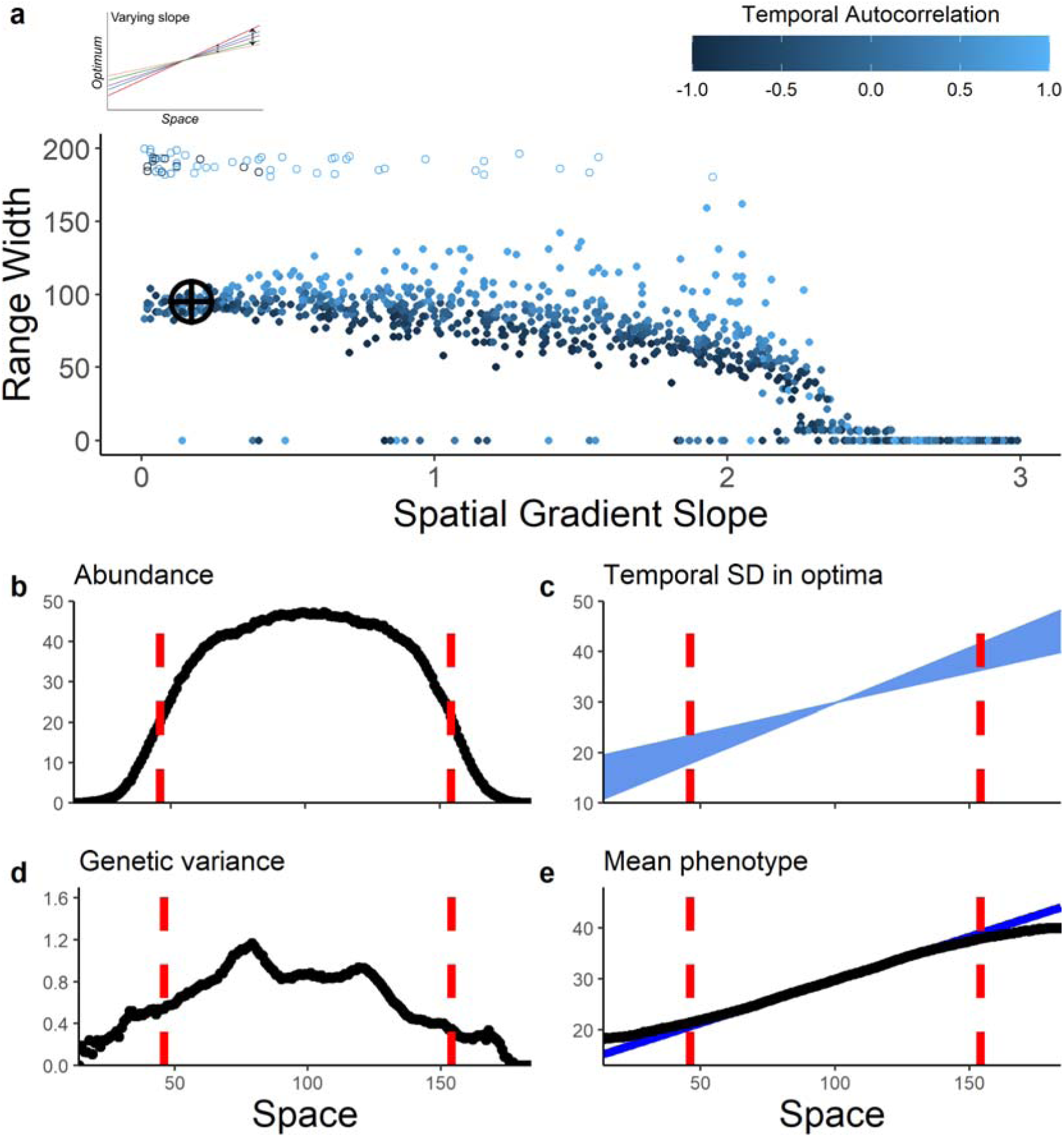
Range dynamics under the “varying slope” scenario. When temporal variance increases away from the landscape center, stable range limits form where temporal variance is too high to allow further adaptation and persistence. (a) Effects of the spatial gradient slope (X-axis) and degree of temporal autocorrelation (point color) on a species’ ultimate range width (Y-axis). (Range width is the number of patches between the species’ two stable range limits.) Each point represents a simulation (N = 1000) at the end of 20,000 generations. Open points near the top of the Y-axis are simulations where the species reached at least one edge of the landscape. (b) abundance, (c) temporal variation in optima, (d) genetic variance, and (e) mean phenotype of populations in a single simulation (*b* = 0.17, *a* = 0.48; marked with crosshairs in panel a) after formation of stable range limits (dashed red lines), averaging over the final 1000 generations of the simulation. We define stable range limits as the most distal patches that did not go extinct for at least 950 of the 1000 generations prior to the end of the simulation. (c) shows the standard deviation of the temporal changes in phenotypic optima, to illustrate how temporal variance increases away from the landscape center. In (e), black points show the mean phenotype and the solid blue line indicates the underlying mean spatial gradient in phenotypic optima.

Range width (distance between range limits) remained relatively constant across a wide range of spatial gradient slopes, but then began to decrease with steeper spatial gradients (Fig. 4a). The high fitness costs of dispersal across steep gradients (due to maladaptation) combined with the negative fitness effects of temporally fluctuating optima to increase extinction risk more quickly as populations moved away from the center. This led to narrower range widths than for species spreading across shallower gradients. When temporal variation was positively autocorrelated, range width generally increased. If that positive autocorrelation was particularly strong (*a* ≈ 1), sometimes the landscape filled completely, especially when the spatial gradient was shallow (open points in Fig. 4). Because environmental conditions at the landscape core were stable, species usually only went extinct with very steep spatial gradients.

The results presented above for the varying intercept and varying slope scenarios were qualitatively robust to changes in carrying capacity (*K*) and dispersal (*m*) (Supp. mat. S1). In general, larger carrying capacities decrease the parameter space where populations go extinct, likely due to lower demographic stochasticity [e.g., 51] and more efficient selection [e.g., 52]. In contrast, higher dispersal lowered the maximum spatial gradient slope where expansion was possible, likely because this introduced a higher mean fitness cost of dispersal [e.g., 14].

When both the slope and the intercept of the spatial gradient varied through time, there was a contraction in the parameter space where range expansion was possible (Supp mat. S4; Figs. S8, S9, S10). In general, species persistence and range expansion required positive temporal autocorrelation in the gradient intercept (i.e., *a*_*intercept*_ > 0). Range width was overall smaller and more variable when the spatial gradient intercept varied along with the slope (compare Figures 4a and S8b).

## Discussion

Our model results show that, for the parameter space explored here, temporal variation in the environment substantially alters our predictions for range dynamics across spatially variable landscapes. When environmental conditions change from generation to generation similarly across the landscape (“varying intercept” scenario), the ultimate fate of species is determined by the interaction between the degree of temporal autocorrelation in environmental fluctuations and the slope of the spatial environmental gradient. As found in previous results [14], species extinction becomes more likely as the slope of the environmental gradient increases. However, positive autocorrelation of temporal variation can allow populations to expand along even very steep gradients. Negative autocorrelation, though, increases demographic fluctuations and increases extinction risk due to fluctuating phenotypic optima resulting in frequent maladaptation and high genetic load. When temporal variation in the environment increases toward the periphery of a landscape (“varying slope” scenario), stable range limits form. Populations persist where temporal variance is modest, but where temporal variance becomes too large, the associated genetic load reduces fitness such that population adaptation and persistence is not possible. The ultimate width of the species’ range is primarily a function of the underlying spatial environmental gradient. When spatial gradients vary in both intercept and slope through time, range expansion overall becomes less likely. Together these results illustrate how temporal variation in the environment can have a pivotal influence on the likelihood of a species colonizing a new landscape and the location of a taxon’s range limit.

When temporal variance has no spatial structure (“varying intercept”), we can clearly delineate the parameter space where colonizing populations go extinct, versus where they expand across the landscape. When the underlying spatial gradient is shallow, expansion can occur across a wide range of temporal correlation patterns, but increasingly positive autocorrelation is required for expansion as the spatial gradient steepens. Environmental noise in nature seems to largely range from random to positively autocorrelated [47]. Indeed, these simulations indicate that if environmental conditions were strongly negatively autocorrelated through time, colonization and expansion would be very rare. Increasing temporal variability is one predicted (and observed) consequence of contemporary climate change [53], and our simulations suggest this increased variability could reduce the probability that populations will be able to successfully track climatic changes via shifting spatial distributions. For example, the upslope colonization process of an alpine plant due to warming could be stymied if temporal variation is augmented by climate change. Our model suggests increased temporal environmental variation due to climate change could further reduce fitness of edge populations, potentially hampering their ability to track or adapt to changing mean conditions. Furthermore, if a species’ current range limit is due in part to increased temporal environmental variation, then models forecasting future distributions built solely using the mean, and not variance, of predicted climate will likely be inadequate.

The biogeographic fact that all species have limited distributions is often at odds with the ability of evolutionary range limit models to produce *stable* range limits. For instance, when Barton [10] adjusted Kirkpatrick and Barton’s foundational work [9] to allow genetic variance to evolve, range expansion was continuous and limits failed to form. Polechová & Barton [14] provided a solution to this conundrum by incorporating genetic drift and demographic stochasticity in their models, but still found that stable limits only formed with nonlinear spatial gradients [see also [15,16]]. Similarly, in our model there are no stable range limits when there is no spatial trend in temporal variance (i.e., varying intercept scenario). However, we do see stable range limits form when there is a spatial gradient in the magnitude of temporal variance — i.e., temporal variance, and the resulting genetic load, increases nonlinearly with distance from the center of the landscape (varying slope scenario) — even across linear spatial gradients. When viewed collectively, our results and those of Polechová & Barton [14], Polechová [15], and Bridle et al. [16] highlight how nonlinear spatiotemporal trends may be key in generating stable range limits. This work indicates we should perhaps expect qualitatively different patterns in how environments change through time and space at the edge versus the core of a species’ range.

Our results suggest that there is a critical threshold at which temporal variation in the environment can stop range expansion and enforce a stable range limit. Do we find evidence for such a pattern in nature? The idea that environments at the edge of species’ ranges tend to be more temporally variable has been assumed more often than empirically shown, but some demographic studies do suggest range edge habitats to be more temporally variable than range core habitats [48,49,54–57]. Beyond indirect inference of temporal environmental variation via demographic data, there are surprisingly few direct measurements of temporal environmental variability across species ranges. One exception is Eckhart et al. [58], who showed that for the California annual plant *Clarkia xantiana* ssp. *xantiana*, interannual variability in precipitation increased toward the subspecies’ eastern range margin. Our model supports the notion that increased variability in precipitation could contribute to the stable range limit observed in this species.

These results lead to several testable predictions. First, does temporal variation in the environment increase toward species range edges? For climatic variables this would be fairly straightforward to test, as we have excellent data for both species distributions and long-term weather. With such data we could also ask whether, looking across species distributions, we see a positive relationship between the steepness of putatively important spatial gradients across a species range and the temporal autocorrelation in that environmental variable. Figure 3 would suggest this relationship — colonization across steep spatial gradients should only be possible in fairly stable environments (i.e., environments with positive temporal correlation). For example, we might expect to observe that species with populations distributed across steep altitudinal gradients experience more positively correlated temporal variation than species spread across more shallow spatial gradients (e.g., a primarily latitudinal rather than altitudinal gradient).

Here we have focused on spatio-temporal environmental variation and its influence on adaptation in populations of a single species. Our understanding of species range dynamics could be further improved by extending this model to include other genetic, life history, and ecological factors that can potentially have large influences on population dynamics and spread in nature. As Antonovics [59] pointed out, evolution of *multiple* traits may often be required for populations to expand into novel habitats. Genetic correlations between traits whose evolution is required to colonize novel habitat may greatly influence the probability of colonization [60,61]. Incorporating phenotypic plasticity and its evolution would also be a valuable extension of the current model [62]. In terms of life history, the effects of overlapping generations may be very important in temporally variable environments [46], especially if an organism’s life cycle includes dormant stages (e.g., seed banks). The addition of species interactions such as competition would further illuminate how ecological phenomena interact with evolutionary processes to modulate range dynamics [12,63,64]. Simulating spread across a patchy landscape, as opposed to the smooth gradients in trait optima modeled here, could also substantially change our predictions for population colonization and spread. Lastly, temporal variation as modeled here is strongly spatially autocorrelated — *i.e*., adjacent patches always experience temporal variation in optima in the same direction, positive or negative. It would be fruitful to relax this assumption to allow less spatial autocorrelation in temporal variation, as the effects of gene flow on adaptation would likely differ.

Species range limits are as ubiquitous as they are puzzling. What prevents adaptation from allowing species distributions to continually grow by “accretion like the rings of a tree” [65]? Recent theory highlights the importance of genetic drift and nonlinear spatial gradients in controlling range dynamics and the location of stable range limits [14–16]. Here we showed how temporal variation in the environment, an ever-present feature of natural systems, strongly contributes to determining whether a population will expand into novel habitat or go extinct, and can readily enforce stable range limits. By expanding range dynamics models to the temporal dimension, we gain more realistic, comprehensive insight into the mechanisms and processes underlying biogeographic patterns, insights of great relevance to invasion biology, the limits to adaptation, and the fate of populations with environmental change.

## Supporting information

Supp. mat.

## Acknowledgements

We thank three anonymous reviewers for their insightful comments that greatly improved the manuscript.

