## Supplementary material for "Increasing temporal variance leads to stable species range limits": Supp. mat.

**Table S1.** Parameter values for the SLiM simulation model.

| Parameter | Definition | Value |
|---|---|---|
| | *Varying parameters* | |
| *b* | Slope of spatial gradient in phenotypic optima | [0-3] |
| *a* | Degree of temporal autocorrelation | [-0.99-1] |
| | *Static parameters* | |
| *K* | Patch carrying capacity (individuals) | 50 |
| *r* | Maximum rate of increase | 0.8 |
| *s* | Strength of stabilizing selection | 0.125 |
| *m* | Expected dispersal per generation (mean of Poisson dispersal kernel) | 0.8 |
| $\mu$ | Mutation rate per base position per generation | $10^{-7}$ |
| *r* | Recombination rate (crossover events per base position per generation) | $10^{-8}$ |
| $\boldsymbol{\tau}_{intercept}$ | Variance of the Gaussian distribution used to implement temporal stochasticity in the “varying intercept” scenario | 4 |
| $\boldsymbol{\tau}_{slope}$ | Varianc of the Gaussian distribution used to implement temporal stochasticity in the “varying slope” scenario | 0.0025 |

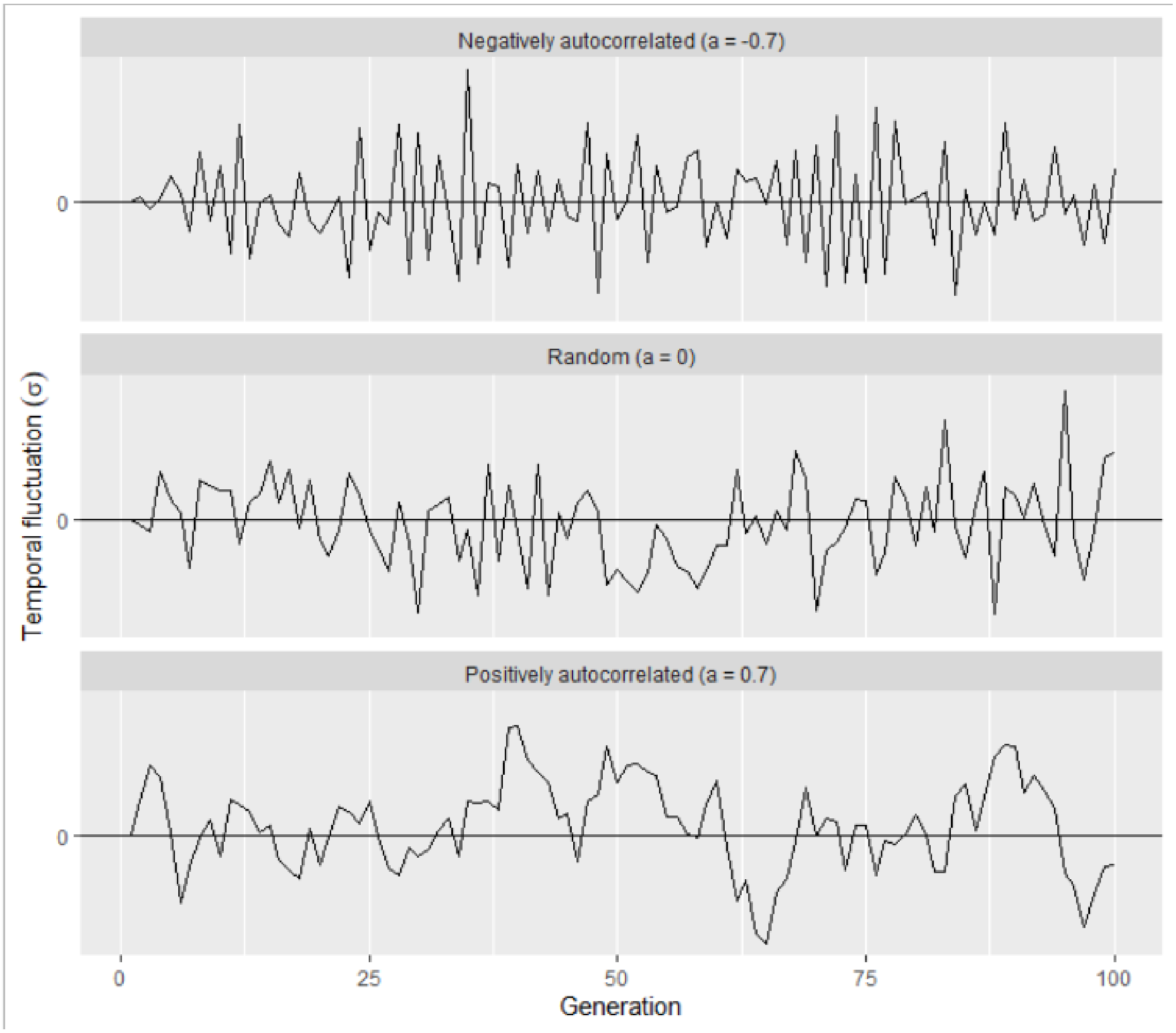


**Figure S1.** Examples of temporal fluctuation patterns in the intercept (for varying intercept scenario) or slope (for the varying slope scenario) of the spatial gradient under negative ($a$ = -0.7), random ($a$ = 0), and positive ($a$ = 0.7) temporal autocorrelation.

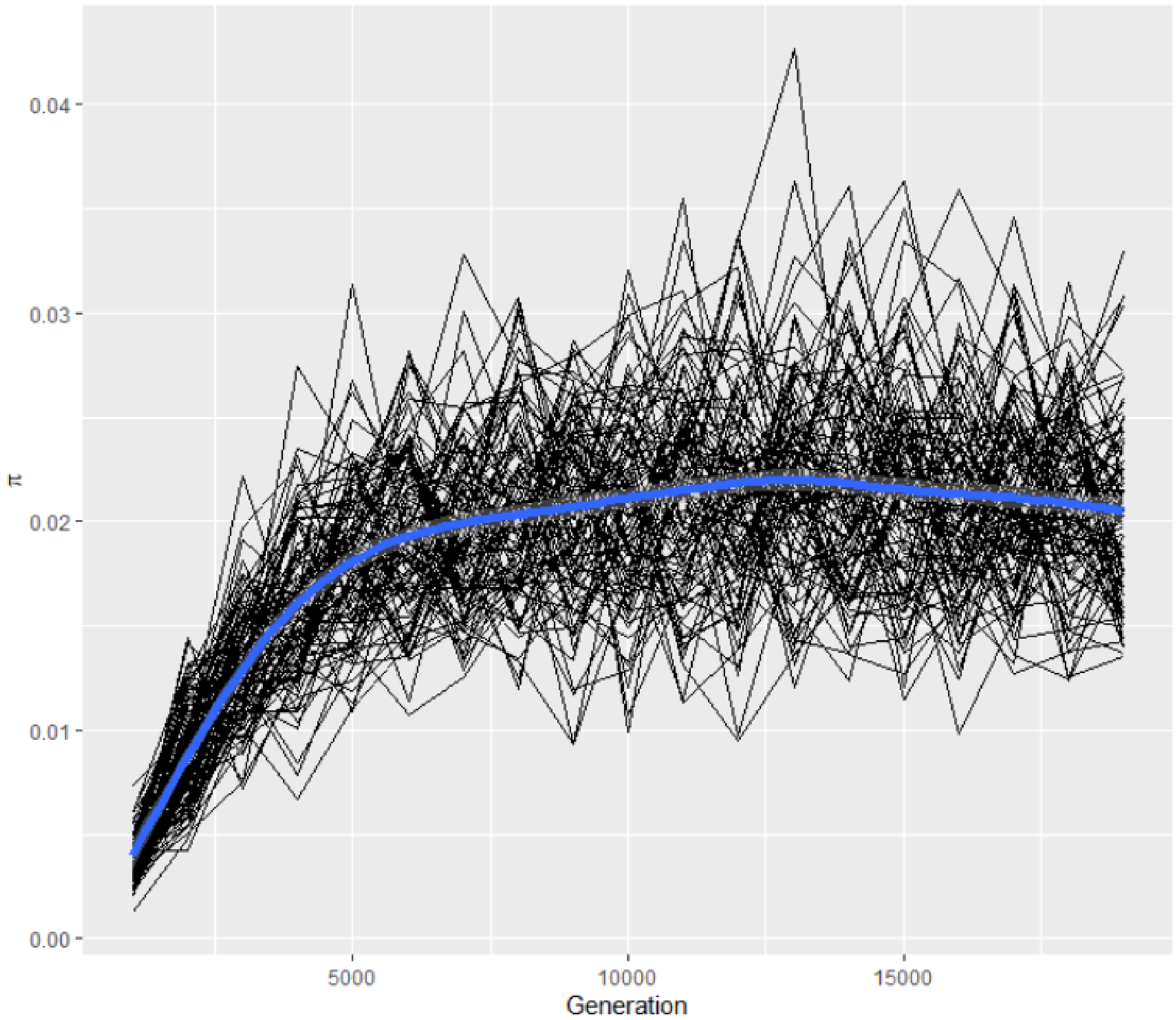


**Figure S2.** Mean heterozygosity of neutral mutations (π) in the central population across time during the 20,000 generation burnin period. 100 simulations are shown (black lines), as well as a loess regression summarizing the trend across all simulations (blue line).

### S1. Sensitivity analyses

#### Carrying capacity

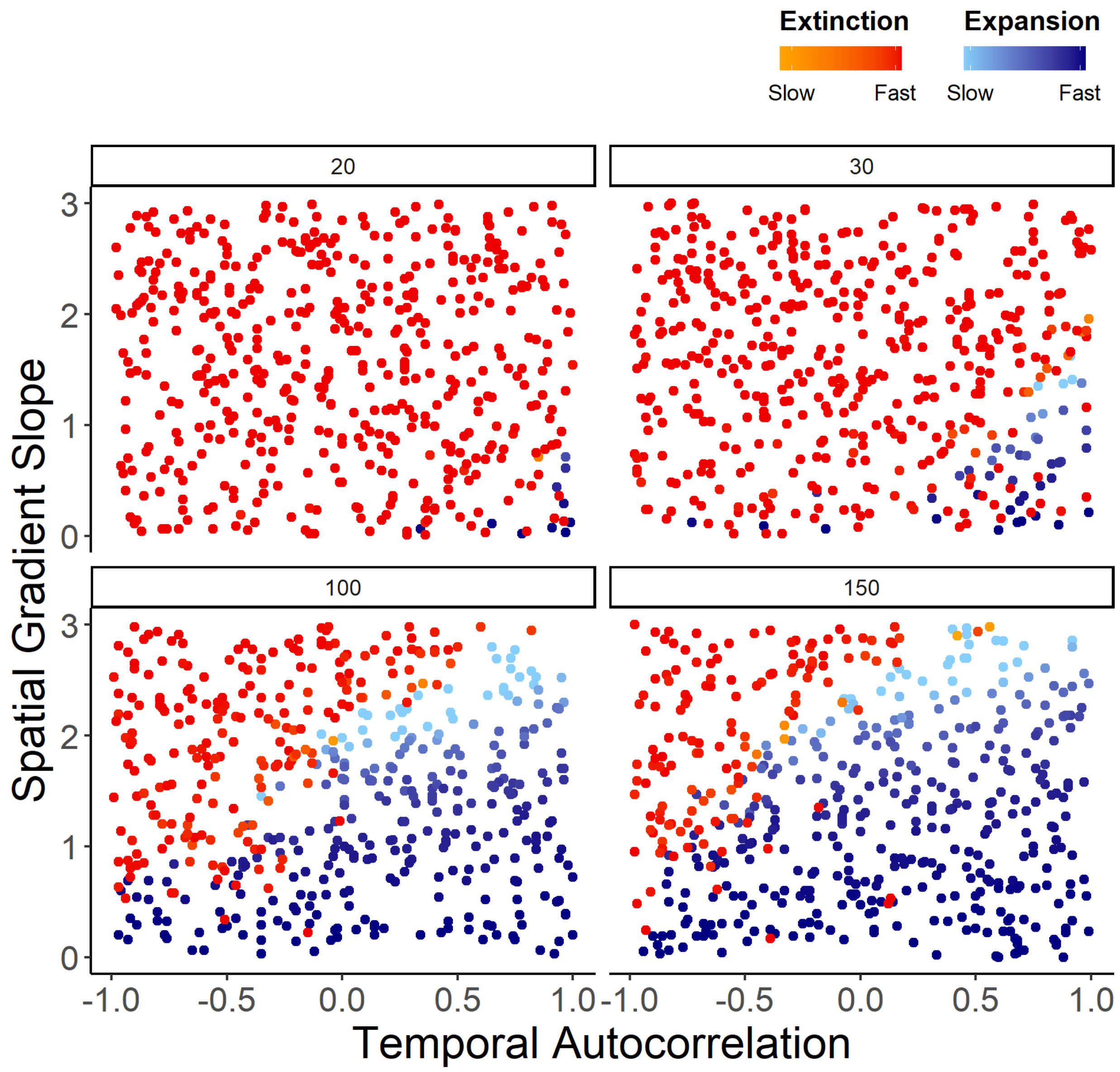


**Figure S3.** Range dynamics under the "varying intercept" scenario for different values of patch carrying capacity ($K = 20, 30, 100, 150$). Compare to Fig. 3, where $K = 50$. Each point represents a simulation (N = 1000). No stable range limits formed in this scenario; species either continually expanded or went extinct. All blue points are simulations where species were able to spread, and red/orange points are simulations where species went extinct; points are colored by how quickly they expanded or went extinct.

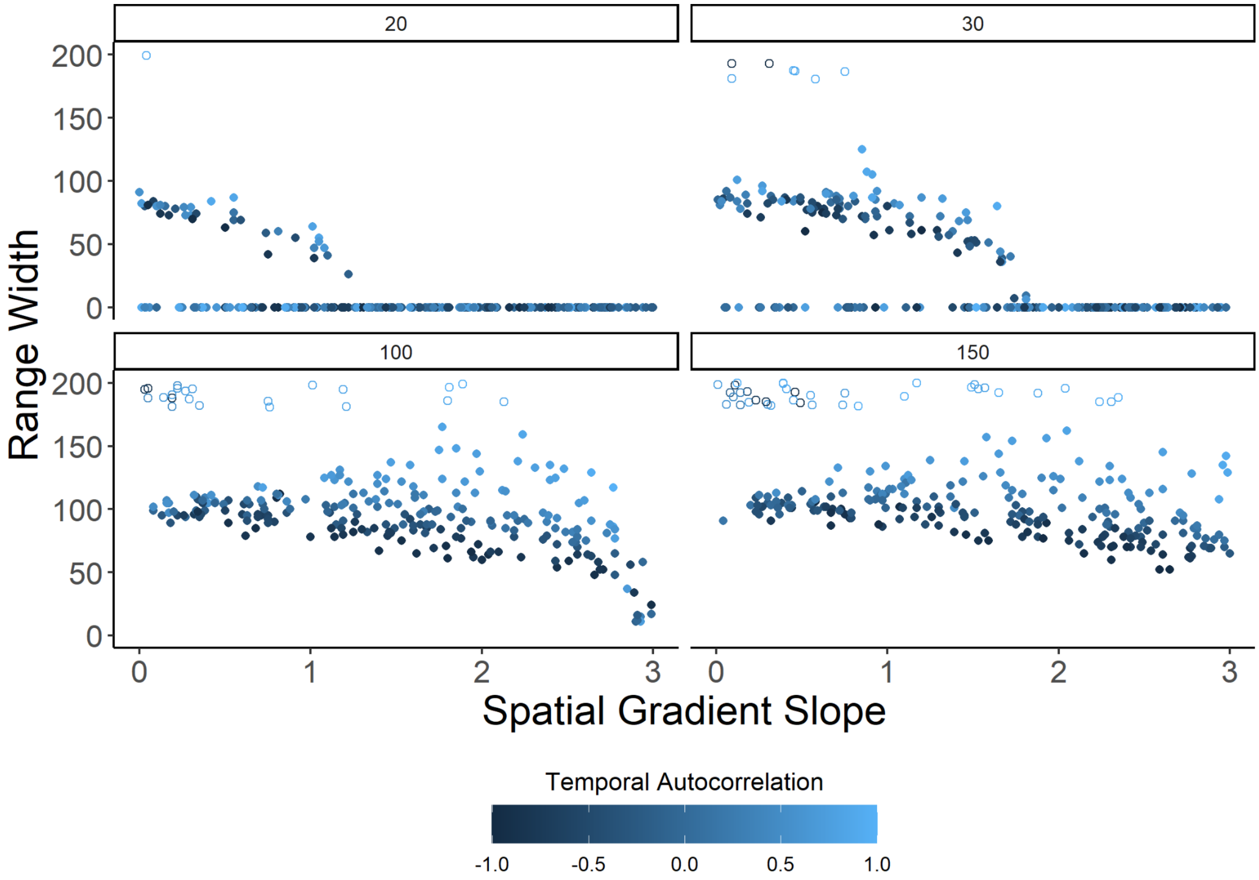


**Figure S4.** Range dynamics under the "varying slope" scenario for different values of patch carrying capacity (*K* = 20, 30, 100, 150). Compare to Fig. 4, where *K* = 50. Plot shows effects of the spatial gradient slope (X-axis) and degree of temporal autocorrelation (point color) on a species' ultimate range width (Y-axis). (Range width is the number of patches between the species' two stable range limits.) Each point represents a simulation at the end of 20,000 generations. Open points near the top of the Y-axis are simulations where the species reached at least one edge of the landscape.

Dispersal rate

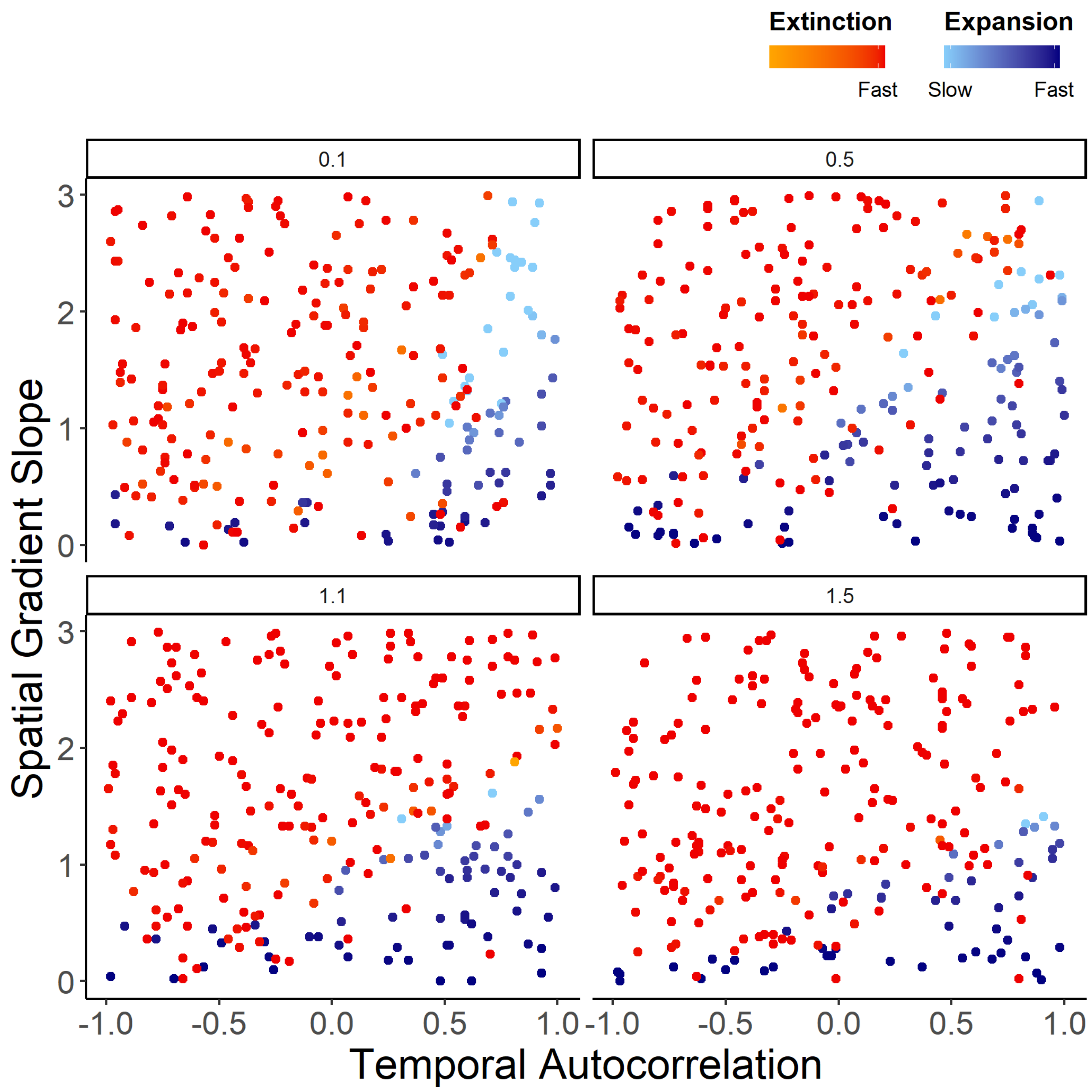


**Figure S5.** Range dynamics under the "varying intercept" scenario for different values of dispersal (*m* = 0.1, 0.5, 1.1, 1.5). Compare to Fig. 3, where *m* = 0.8. Each point represents a simulation. No stable range limits formed in this scenario; species either continually expanded or went extinct. All blue points are simulations where species were able to spread, and red/orange points are simulations where species went extinct; points are colored by how quickly they expanded or went extinct.

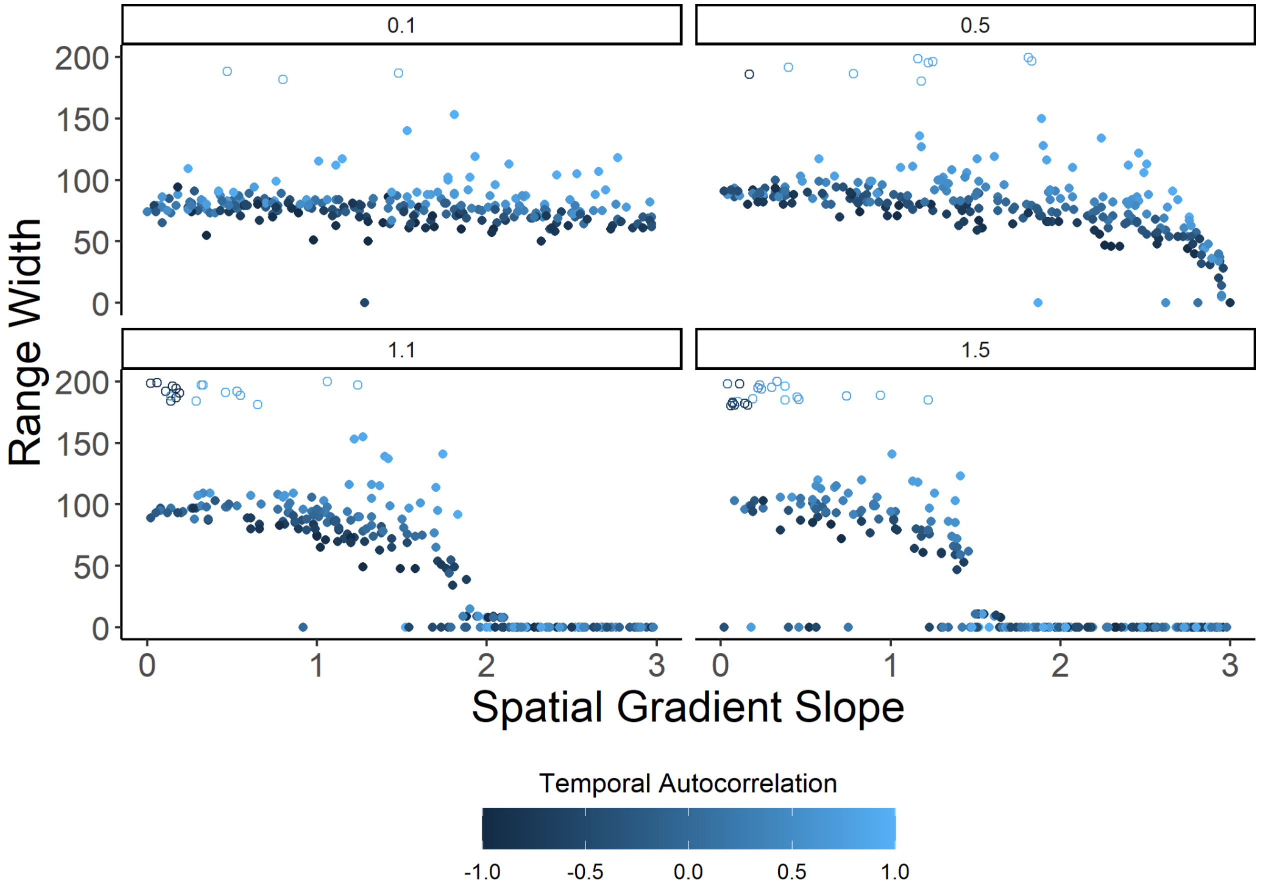


**Figure S6.** Range dynamics under the “varying slope” scenario for different values of dispersal ($m$ = 0.1, 0.5, 1.1, 1.5). Compare to Fig. 4, where $m$ = 0.8. Plot shows effects of the spatial gradient slope (X-axis) and degree of temporal autocorrelation (point color) on a species’ ultimate range width (Y-axis). (Range width is the number of patches between the species’ two stable range limits.) Each point represents a simulation at the end of 20,000 generations. Open points near the top of the Y-axis are simulations where the species reached at least one edge of the landscape.

### S2. Selection gradients

In natural populations, selection on traits is often measured using standardized linear selection gradients, β, which describe how an individual’s fitness varies with its trait value (Lande and Arnold 1983). The magnitude of temporal fluctuations in these gradients can be described with $\sigma_\beta$ (e.g., de Villemereuil et al. 2020). To compare the fluctuations in selection that emerge in our models to what is observed in natural populations, we estimated β and $\sigma_\beta$ given the parameter values used in our simulations. We first generated 100,000 draws from $\Phi_t \sim$ Normal(0, 4); these values represent deviations from the long-term mean optimum across 100,000 generations, assuming no temporal autocorrelation (i.e., $a$ = 0). Following (Lande and

Arnold 1983), linear selection gradients were estimated for each generation as the slope coefficient of the linear regression of mean-relativized fitness on the phenotypic trait values of two hypothetical phenotypes: (1) a phenotype perfectly adapted to the current trait optimum and (2) a phenotype adapted to the long-term mean optimum. Per (Lande and Arnold 1983), we standardized this gradient by multiplying it by the phenotypic standard deviation, $\sigma_P$, which we estimate at 1.8 (the mean $\sigma_P$ in the central patch in our "varying intercept" simulations after 20,000 generations). With this simulation of temporal stochasticity across 100,000 generations, the absolute value of the standardized linear selection gradient, $|\beta|$, averaged ~0.10 across generations, and $\sigma_\beta \approx 0.14$. These selection gradients are well within the range of selection gradients and their variance in nature, [median $|\beta| = 0.16$ in Kingsolver et al. (2001); $\sigma_\beta = 0.099$ in de Villemeruil et al. (2020), averaging across birds and mammals]. In the varying slope scenario, $|\beta|$ and $\sigma_\beta$ increased with distance from the landscape center as temporal variance increased, matching temporal variance of the varying intercept scenario 40 patches away from the landscape center (Fig. 2).

#### S3. Comparison with Polechová and Barton 2015

We compared the results of our simulation model (with no temporal stochasticity; i.e., $a = 1$) with those predicted by the model used in Polechová and Barton 2015. Their model showed that range expansion was prevented at a "threshold" environmental gradient,

$$B \gtrsim 0.15 N\sigma\sqrt{s}$$

where $B$ is the effective environmental gradient, $N$ is the size of each deme/patch, σ is the SD of a discretized and truncated Gaussian dispersal kernel, and $s$ is the strength of selection.

The effective environmental gradient, $B$, is

$$B = b\sigma / r^{\star}\sqrt{2V_s}$$

where $b$ is the spatial gradient in the environmental optimum, $r^{\star}$ is the rate of return to equilibrium population, and $V_s$ is the variance of stabilizing selection ($V_s = 1/2s$). $r^{\star}$ is defined as

$$r^{\star} = r_m - V_G / 2V_s$$

where $r_m$ is the maximum exponential growth rate and $V_G$ is genetic variance, $V_G = b\sigma\sqrt{V_s}$.

Because our model builds from Polechová and Barton 2015 and the related Bridle et al. 2019, most of these parameters map directly to our model. The only significantly different parameter is dispersal, σ. Our dispersal parameter, $m$, is the mean of a Poisson dispersal kernel, as opposed to σ, which is the SD of a truncated, discretized Gaussian dispersal kernel. For the purposes of comparing our model to Polechová and Barton 2015, we estimate an $m = 0.8$, as used in our model, is approximately equal to a $\sigma = 1.1$. We can then estimate the threshold $B$ in our model using the following parameter values:

| **Parameter** | **Value** |
|---|---|
| $b$ | *Varies* |
| σ | 1.1 |
| $N$ | 50 |
| $s$ | 0.125 |
| $V_s$ | 4 |
| $r^\star$ | *Varies with b* |
| $r_m$ | 0.8 |
| $V_G$ | *Varies with b* |

Using these parameter values in the equations above, the threshold $B$ in our model is predicted to occur at $b \approx 1.96$. The observed threshold from our simulations was $b \approx 2.5$ (Fig. SX). The discrepancy likely occurs due to differences in mating system (we are modeling sexual diploids, while Polechová and Barton 2015 model haploids) and dispersal (we are modeling dispersal with a Poisson kernel, as opposed to a Gaussian distribution).

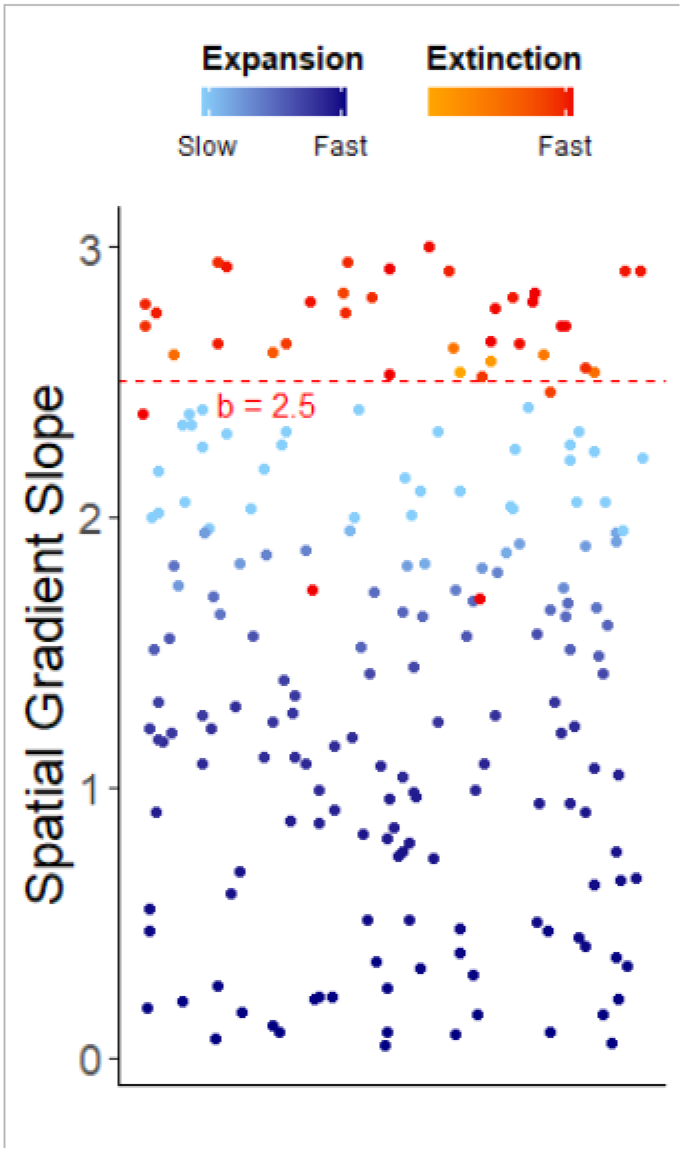


**Fig. S7.** Fate of populations with a temporally constant environment ($a$ = 1). No variable is represented along the X axis but points are jittered horizontally for clarity.

#### S4. Varying intercept and slope

When both the slope and the intercept of the spatial gradient varied through time, there was a contraction in the parameter space where range expansion was possible (Fig. S2). In general, species persistence and range expansion required positive temporal autocorrelation in the gradient intercept (i.e., $a_{intercept} > 0$). The degree of autocorrelation in slope of the gradient had only a modest influence on extinction and range width (Figs. S3, S4), and so here we focus on scenarios with no autocorrelation in gradient slope (i.e., $a_{slope} = 0$) for simplicity. Temporal variation in the slope (regardless of the degree of autocorrelation) increased extinction compared to scenarios with only a varying intercept (compare Figures 3A and S2A). Interestingly, in the parameter space where temporal autocorrelation in gradient intercept was positive and thus expansion was possible, simulations in which the slope of the spatial gradient was very shallow were more likely to go extinct than those with steeper spatial gradients (Fig. S2A, lower right corner). This is because steeper spatial gradients increased genetic variance across the landscape via dispersal (Fig. S7), which better equipped populations to withstand temporal fluctuations in optima. Due to varying gradient slopes (and thus an increase in temporal variance with distance from the landscape center), stable range limits formed for all species that avoided extinction (no extant species expanded to fill the landscape). Range width was overall smaller and more variable when the spatial gradient intercept varied along with the slope (compare Figures 4A and S2B).

### Other supplementary figures

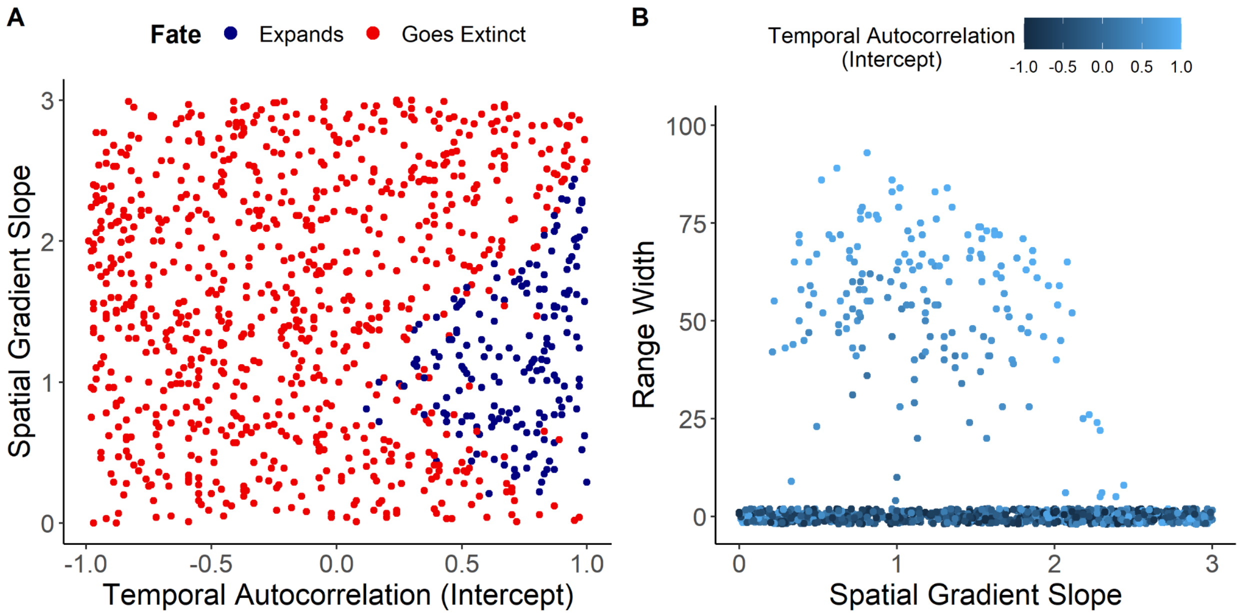


**Figure S8.** Range dynamics when both the slope and intercept of the spatial gradient vary through time. Each point represents a simulation (N = 1000). (A) shows the fate of species (expansion vs. extinction); compare to Fig. 2. All blue points are simulations where species were able to spread, and red points are simulations where species went extinct. (B) shows effects of the spatial gradient slope and degree of temporal autocorrelation in gradient intercept (point color) on a species' range width after the formation of stable range limits; compare to Fig. 3 (but note different Y-axis limits). Species with range width = 0 (jittered points at bottom of plot) went extinct; stable range limits formed for species that avoided extinction. In both (A) and (B), the degree of temporal autocorrelation in gradient slope is fixed at $a = 0$ (i.e., random fluctuations in slope). Three dimensional plots including a range of $a$ values for the varying slope parameter are found in Figures S2 and S3.

**Figure S9.** (View file **figS9.html**.) Fate of populations when both the slope and intercept of the spatial gradient vary through time. Each point is a simulation (N = 1000). This plot includes a range of autocorrelation values (*a*) for slopes, as opposed to Fig. S8A where $a_{slope}$ was fixed at zero. All blue points are simulations where populations were able to spread, and red points are simulations that went extinct.

**Figure S10.** (View file **figS10.html**.) Range width when both the slope and intercept of the spatial gradient vary through time. Each point is a simulation (N = 1000). This plot includes a range of autocorrelation values (*a*) for slopes, as opposed to Fig. S8B where $a_{slope}$ was fixed at zero. Plot shows effects of the spatial gradient slope, degree of temporal autocorrelation in gradient intercept, and degree of temporal autocorrelation in gradient slope (point color) on a species' range width after the formation of stable range limits. Simulations with range width = 0 went extinct and are marked with hollow points.

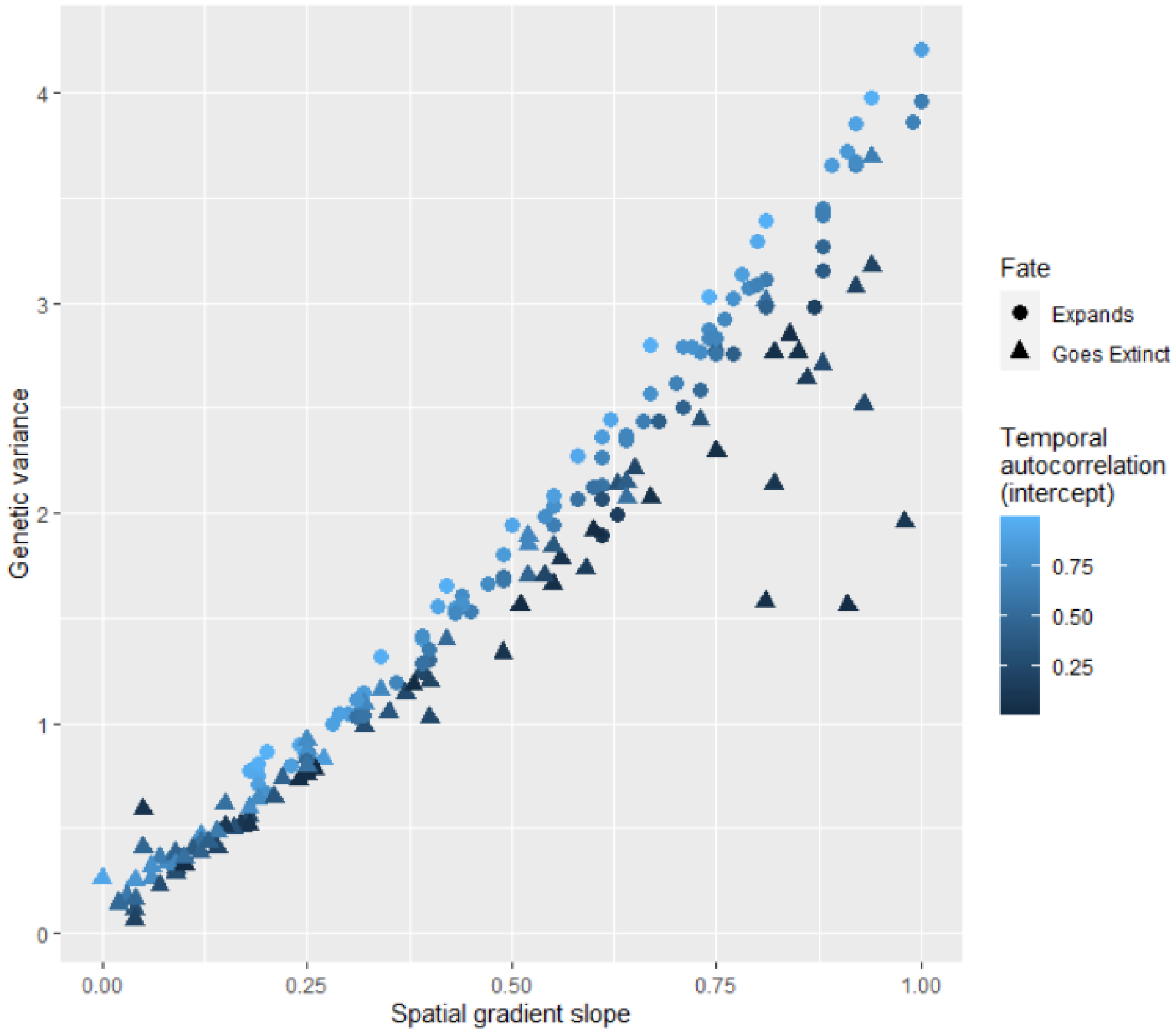


**Figure S11.** Genetic variance for the trait conferring adaptation to the environmental gradient increases as the spatial gradient slope steepens in the varying intercept / random slope scenario (Fig. S8).

### References

de Villemereuil, P., A. Charmantier, D. Arlt, P. Bize, P. Brekke, L. Brouwer, A. Cockburn, S. D. Côté, F. S. Dobson, S. R. Evans, M. Festa-Bianchet, M. Gamelon, S. Hamel, J. Hegelbach, K. Jerstad, B. Kempenaers, L. E. B. Kruuk, J. Kumpula, T. Kvalnes, A. G. McAdam, S. E. McFarlane, M. B. Morrissey, T. Pärt, J. M. Pemberton, A. Qvarnström, O. W. Røstad, J. Schroeder, J. C. Senar, B. C. Sheldon, M. van de Pol, M. E. Visser, N. T. Wheelwright, J. Tufto, and L.-M. Chevin. 2020. Fluctuating optimum and temporally variable selection on breeding date in birds and mammals. Proc. Natl. Acad. Sci. U. S. A. 117:31969–31978.

Kingsolver, J. G., H. E. Hoekstra, J. M. Hoekstra, D. Berrigan, S. N. Vignieri, C. E. Hill, A. Hoang, P. Gibert, and P. Beerli. 2001. The strength of phenotypic selection in natural populations. Am. Nat. 157:245–261. journals.uchicago.edu.

Lande, R., and S. J. Arnold. 1983. THE MEASUREMENT OF SELECTION ON CORRELATED CHARACTERS. Evolution 37:1210–1226. Wiley Online Library.
